# speciesgeocodeR: An R package for linking species occurrences, user-defined regions and phylogenetic trees for biogeography, ecology and evolution

**DOI:** 10.1101/032755

**Authors:** Alexander Zizka, Alexandre Antonelli

**Affiliations:** University of Gothenburg, Department of Biological and Environmental Sciences, Carl Skottsbergs gata 22B, P.O. Box 461, SE 405 30, Göteborg, Sweden; Gothenburg Botanical Garden, Department of Biological and Environmental Sciences, Carl Skottsbergs gata 22A, SE 41319, Göteborg, Sweden

**Keywords:** Geographic information system (GIS), biodiversity, data quality

## Abstract

1. Large-scale species occurrence data from geo-referenced observations and collected specimens are crucial for analyses in ecology, evolution and biogeography. Despite the rapidly growing availability of such data, their use in evolutionary analyses is often hampered by tedious manual classification of point occurrences into operational areas, leading to a lack of reproducibility and concerns regarding data quality.

2. Here we present speciesgeocodeR, a user-friendly R-package for data cleaning, data exploration and data visualization of species point occurrences using discrete operational areas, and linking them to analyses invoking phylogenetic trees.

3. The three core functions of the package are 1) automated and reproducible data cleaning, 2) rapid and reproducible classification of point occurrences into discrete operational areas in an adequate format for subsequent biogeographic analyses, and 3) a comprehensive summary and visualization of species distributions to explore large datasets and ensure data quality. In addition, speciesgeocodeR facilitates the access and analysis of publicly available species occurrence data, widely used operational areas and elevation ranges. Other functionalities include the implementation of minimum occurrence thresholds and the visualization of coexistence patterns and range sizes. SpeciesgeocodeR accompanies a richly illustrated and easy-to-follow tutorial and help functions.

## Introduction

Species distributions and phylogenetic trees constitute core data in biogeography, ecology and evolution. Ancestral range estimation and area-specific diversification rate analyses are two prominent examples (e.g. Meseguer et al., 2014; Antonelli et al., 2015). Most methods applied in this context depend on the classification of taxonomic occurrences (typically species or populations) into discrete geographic units, such as continents, biomes, mountain ranges, or user-defined operational areas. These methods include, among others, Lagrange (Ree & Smith 2008), BayesRates (Silvestro et al. 2011), GeoSSE/diversitree (Goldberg et al. 2011; FitzJohn 2012), BAT (Cardoso et al. 2015) and BioGeoBEARS (Matzke 2013). However, detailed information on species distributions is often available as GPS-based point locations, for example from collected specimens or field observations. A “manual” classification of point occurrences into operational areas using expert knowledge or graphical user interfaces, including GIS software, is feasible for small-scale studies. However, it becomes time consuming and error-prone as the number of species and areas increase, or when the spatial resolution for the classification increases (e.g., assigning species to environmentally and topographically complex regions). Manual data curation is particularly impractical for large-scale analyses using data from public sources, such as the Global Biodiversity Information Facility (GBIF, www.gbif.org), ebird (Sullivan et al. 2009) or SUPERSMART (Antonelli et al. 2014) which has led to a discussion on data-quality and reproducibility of large scale studies (e.g. Yang et al., 2013; Hjarding et al., 2014; Maldonado et al., 2015).

Here we present speciesgeocodeR, an R package to automatically clean, process and analyse species occurrence data and to code them into discrete areas. The core functionalities of the package are: 1) automated and reproducible data cleaning, 2) the rapid and reproducible classification of point occurrences into discrete operational areas, in an adequate format for subsequent phylogenetic analyses, including widely used nexus files as well as input files for the BioGeoBEARS package (Matzke 2013) and 3) a publication-quality and rich visualization of the results. The package summarizes occurrence and species numbers per area, which allows the rapid exploration of even very large datasets, and provides additional functionalities such as: including elevation ranges and occurrence thresholds in the area classification, automatic downloading of GBIF data and WWF biomes and ecoregions (Olson et al. 2001), as well as calculating and visualizing coexistence patterns and species ranges. SpeciesgeocodeR is a tool for exploration, visualization and quality control of species distribution. The package is particularly suitable for (but not restricted to) the preparation of input files for use with phylogenetic trees, such as for biogeographic reconstructions or diversification rate analyses.

## Description

SpeciesgeocodeR is written for the use in R (R Developement Core Team 2015). It can handle datasets of any size (from a few species to hundreds of thousands), and is particularly user-friendly, also for R-beginners. A comprehensive tutorial (“Data cleaning and exploration with speciesgeocodeR”) is provided as part of the package (Supplementary file 1). SpeciesgeocodeR takes advantage of methods and functionalities developed in the sp (Pebesma & Bivand 2005), raster (Hijmans 2014b), maps (Becker et al. 2013) and maptools (Bivand & Lewin-Koh 2013) packages. Furthermore some particular functionalities use the mapdata (Becker et al. 2014), rgbif (Chamberlain et al. 2015) and geosphere (Hijmans 2014a) packages.

## Workflow example and case study

We demonstrate a typical workflow with the major functions of the package on a case study of Lemur *(Lemuriformes)* distributions in Madagascan biomes. Figure 1 shows the results and mentions the functions applied. For illustration we use occurrences from GBIF (http://data.gbif.org 2015) and a simplified version of the WWF biomes for Madagascar as example data. The occurrence dataset included global records, and also comprised individuals in captivity. As lemurs in the wild are restricted to Madagascar, we use these specimens to illustrate one function of the automated data cleaning. The data are distributed together with the package. The code to reproduce the analyses can be found in the tutorial (Supplementary file 1) together with additional examples.

**Figure 1.**
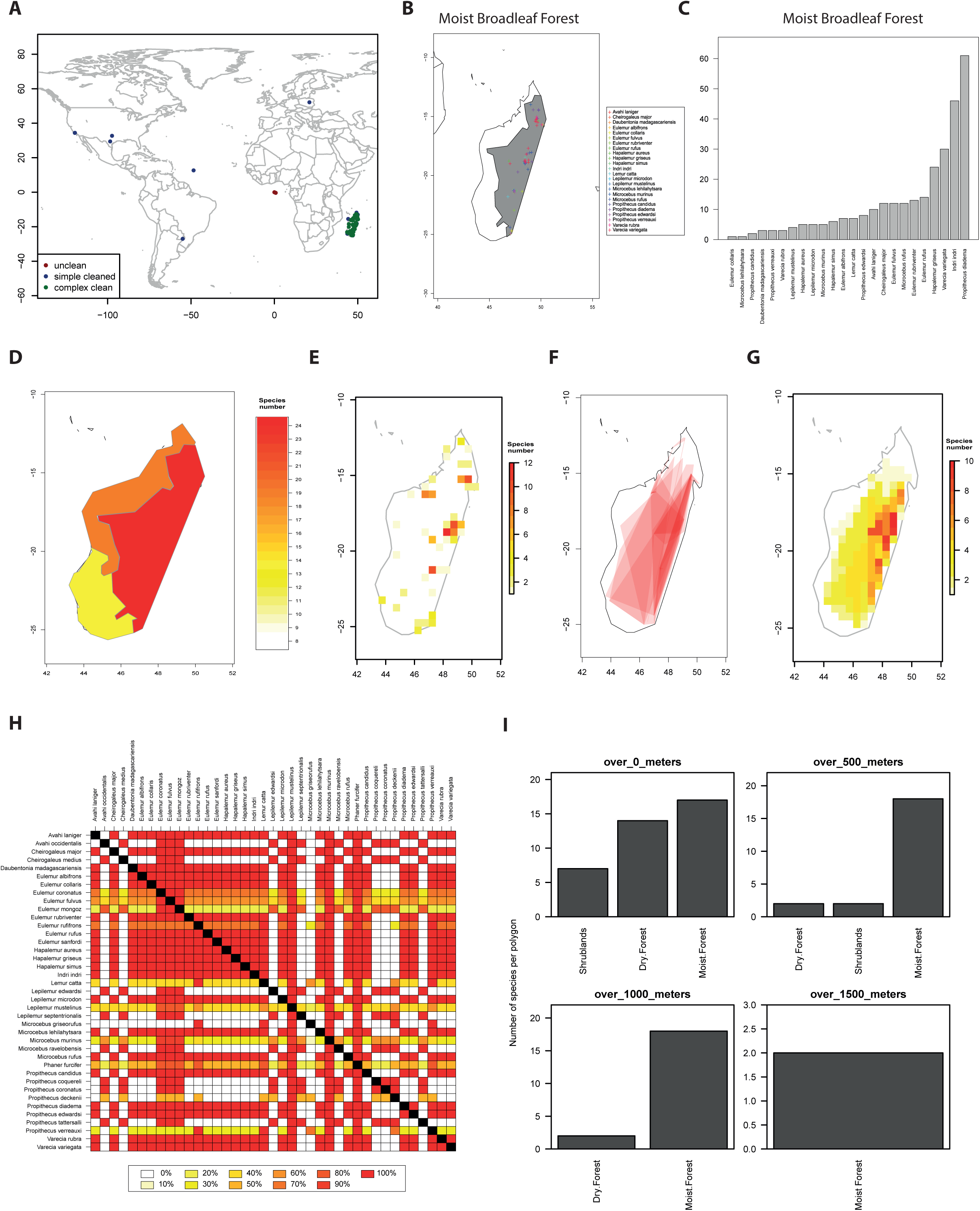
The major functions of the speciesgeocodeR package (function name in *italics)* demonstrated for lemurs. A) Results of the automated cleaning of geographic occurrence data *(GeoClean)*, red = records flagged by checks for coordinate validity, blue = records flagged by test for country borders. B) Lemur species occurring in moist broadleaf forest, as an example of the summary maps automatically produced by *SpGeoCod*. C) The number of occurrence records for each species in Moist Broadleaf Forest *(SpGeoCod)*. D) Lemur species richness in the biomes of Madagascar *(MapRichness)*. E) Species richness raster of lemur species in Madagascar *(RichnessGrid, MapGrid)*. F) Overlaid distribution Ranges of 26 Lemur species *(CalcRange, PlotHull)*. G) Species richness raster derived from the ranges shown in I) *(RangeRichness)*. H) Heatplot of the coexistence matrix of 39 Lemur species in three Madagascan biomes. The colors indicate the percentage (per row) of shared occurrences given the biomes *(CoExClass)*. I) Species number in different elevation ranges in each biome *(SpGeoCod(elevation = T, threshold = c(500,100,1500))*

### DATA CLEANING AND DATA QUALITY

The quality of occurrence data downloaded from public databases has often been debated and, less often, tested (e.g. Yang et al., 2014; Maldonado et al., 2015; Meyer et al., 2015). With the *GeoClean* function speciesgeocodeR offers an automated and reproducible flagging of potentially problematic records (see tutorial for code examples). The function includes basic tests for coordinate validity (e.g. invalid coordinates, zero coordinates, equal latitude and longitude) and more complex tests accounting for common problems in large datasets (e.g. if occurrences fall within the right country borders, if occurrences have been assigned to the country centroid or country capital or to the GBIF headquarters). Depending on the “verbose” argument (determining the amount of output information), the result can be a single vector summarizing the results of all tests, or a “data.frame” table with results of each individual test. Figure 1A shows the results of automated cleaning of the example dataset. Of 627 records, 224 were flagged as problematic (red and blue on the map) and excluded from subsequent analyses.

### OCCURRENCE CLASSIFICATION

The core function of the package, *SpGeoCod*, classifies species occurrences into operational areas and calculates summary tables. The standards input are sets of (1) species occurrence points and (2) geographical areas (polygons). Both can be provided as R objects of the classes “data.frame” and “SpatialPolygons”, or as tab delimited .txt and .shp files in the working directory. Alternatively, a vector of species names can be used as occurrence data. In this case the occurrences will be downloaded from GBIF. In a similar manner, the WWF ecoregion and biomes (Olson et al. 2001) can be directly included as areas via the *WwfLoad* function. *SpGeoCod* offers a set of options to customize the area classification. For example, it is possible to condition the presence of a species in a given area on a minimum number of occurrences (as percentage of total species occurrences), or to include user-defined elevation zones into the classification. If desired, the entire analysis can be run with a single line of code after the data is uploaded:

*data(lemurs)*
*data(mdg_ biomes)*
*outp <- SpGeoCod(lemurs, mdg_poly, areanames = “name”)*

The results of *SpGeoCod* are saved in an object of the S3 class “spgeoOUT”, which can be explored using plot or summary methods. The *WriteOut* function exports the area classification in nexus format for the direct use with phylogenetic software or as a text file in the adequate format for the BioGeoBEARS package. Additionally the function can create a set of graphics and maps (Fig 1A-I) in pdf format, and summary tables in text format. As part of the typical output, figure 1B shows an overview map of all lemur occurrences in broadleaved tropical forests (the most species rich biome in Madagascar) and figure 1C shows the occurrence number per species in this biome. Similar statistics and maps are automatically created for each area and each species, and could e.g. be included as supplemental files in publications.

### DATA EXPLORATION

SpeciesgeocodeR includes multiple additional functions to explore the data based on the structure of the “spgeoOUT” class (see function help files for details on calculations). Species numbers in each area can be visualized using *MapRichness* (Fig 1D). Grids with species and occurrence numbers can be calculated using *RichnessGrid* and visualized using *MapGrid* (Fig. 1E). Species ranges and range sizes given the occurrence points can be calculated using *CalcRange* and visualized using *PlotHull* (Fig. 1F, mean range size per lemur species = 56,000 km^2^, median = 12,000 km^2^). As for Madagascan lemurs, occurrence data is often scarce and gridded diversity maps can be strongly influenced by sampling patterns (Fig 1E). To circumvent this problem *RangeRichness* can create richness maps based on the species ranges (Fig 1G). A coexistence matrix of the species in the given polygons can be calculated using *CoExClass* and visualized as a heat plot using *plot* (Fig 1H). Figure 1I shows an example of including user-defined elevation zones in the analysis, showing the species number per area and elevation range, produced with *SpGeoCod(elevation = T, threshold = c(500, 100, 1500)*. Elevation is automatically calculated from georeferences by linking them to the CGIAR-CSI SRTM 90m digital elevation data (http://srtm.csi.cgiar.org/) that cover the world at 90m resolution.

## Comparison with other tools

SpeciesgeocodeR adds new functionalities to toolbox of recent, primarily macroecological packages in the R-environment (e.g. modestR, García-Roselló et al., 2014 and letsR, Vilela & Villalobos, 2015). Conceptually, it differs from other packages by its focus on data quality control and the link to evolutionary and biogeographic analyses. Some of the implemented differences between SpeciesgeocodeR and available tools include 1) automated and reproducible data-cleaning, 2) the easy and quick area classification of species into pre-defined polygons producing a format suitable for analyses with phylogenetic trees, 3) the possibility to calculate elevation profiles and code species into elevation zones, 4) allowing the user to specify minimum occurrence thresholds for area classifications, 5) the seamless inclusion of GBIF data and WWF biomes and ecoregions and 6) the calculation of coexistence patterns in the areas of interest. Finally, a particular strength of speciesgeocodeR is the easy analysis and visualization of even very large datasets (including millions of records) with just a few lines of code. In the future, speciesgeocodeR could be easily expanded to carry out additional analyses, including implementations of ancestral state reconstructions and geographic mapping of sister clades.

## Availability

SpeciesgeocodeR is available from CRAN since October 2015 (http://CRAN.R-project.org/package=speciesgeocodeR) under a GPL-3 license, and a developmental version is available from GitHub (https://github.com/azizka/Speciesgeocoder-Rpackage). Users are welcome to join a users’ list (speciesgeoco deR@googlegroups) for receiving updates, reporting issues and providing feedback and suggestions. A python version of speciesgeocodeR is currently being developed ( Töpel et al. 2014).

## Acknowledgments

The authors would like to thank Mats Töpel, Maria Fernanda Calió and Ruud Scharn for discussions and ideas on this project, Esther Nieto and Diogo Provete for testing the package and Angela Cano, Gaëlle Bocksberger and Daniele Silvestro for helpful comments on the manuscript. Funding for this work was provided by the Swedish Research Council (B0569601), the European Research Council under the European Union’s Seventh Framework Programme (FP/2007-2013, ERC Grant Agreement n. 331024), and a Wallenberg Academy Fellowship to A.A.

## Supporting information

Additional Supporting Information may be found in the online version.

**Appendix S1**. SpeciesgeocodeR tutorial

